# Temporal Dynamics and Adaptive Mechanisms of Microbial Communities: Divergent Responses and Network Interactions

**DOI:** 10.1101/2024.08.22.609127

**Authors:** Shengjie Sun, Zhiyi Qiao, Denis V. Tikhonenkov, Yingchun Gong, Hua Li, Renhui Li, Kexin Sun, Da Huo

**Author notes:** **Corresponding author:** Da Huo. These authors contributed equally to this work (Shengjie Sun, Zhiyi Qiao).

## Abstract

Microbial communities are integral to aquatic ecosystems, playing key roles in biogeochemical cycles, nutrient recycling, and ecosystem functioning. Despite their importance, the temporal dynamics of these communities, especially in response to short-term environmental changes, remain inadequately understood. This study utilizes high-throughput amplicon sequencing and Co-occurrence network analysis to elucidate how prokaryotic and eukaryotic microbial communities dynamically respond to short-term environmental fluctuations, and dynamics of networks. Our results reveal distinct temporal patterns, with eukaryotic communities showing consistent increases in diversity, while prokaryotic communities exhibited more pronounced fluctuations and directional changes. Co-occurrence network analysis further highlighted the differences in community structure between phases of homogenization and heterogenization. Specifically, we observed a shift from a more modular network during the homogenization phase to a more centralized network in the heterogenization phase, resulting in increased network fragility. Environmental factors, such as nutrient levels and climatic conditions, were found to play pivotal roles in shaping these dynamics. This study provides critical insights into the resilience and adaptability of microbial communities, emphasizing the intricate interplay between biodiversity, network structure, and ecosystem stability under varying environmental conditions.

## 1 Introduction

Microbial communities play a fundamental role in aquatic ecosystems, serving as essential contributors to biogeochemical cycles, nutrient recycling, and overall ecosystem functioning. Amidst the ongoing challenges posed by global warming and anthropogenic pressures, the vulnerability of microbial communities has increasingly come to the forefront. Due to their small size, rapid growth rates, and metabolic flexibility, microbial communities can exhibit rapid turnover, often within hours to days, allowing them to respond swiftly even to minor disturbances (Faust et al., 2015). These short-term responses often result in significant variations in community dynamics, with microbial abundances fluctuating notably between measurement time points (Gonze et al., 2018). Recent studies have demonstrated that, in marine eukaryotic communities, differences in community composition over a few days were often greater than those observed over several kilometers (Lie et al., 2013). Similarly, short-term dynamics of microbial communities in estuarine and lacustrine systems have shown rapid changes in community composition within 1-3 weeks, indicating a continuous reassembly of these communities (Mangot et al., 2013; Vigil et al., 2009). Recent research suggests that short-term community dynamics may have profound and lasting impacts on both community structure and function (Bloxham et al., 2024). Therefore, understanding the temporal dynamics of these microbial communities is crucial for predicting the resilience and adaptability of aquatic ecosystems under changing environmental conditions.

Biotic factors are equally crucial in driving microbial community dynamics (Fuhrman et al., 2015). As key components of microbial communities, the coexistence of prokaryotic and eukaryotic communities, with their distinct diversity and abundance patterns, plays a pivotal role in shaping microbial community structure and function through their interactions. Research has demonstrated that prokaryotic and eukaryotic communities exhibit distinct temporal variations. For instance, a study conducted in a freshwater lake in the subtropical region of eastern China found that prokaryotic communities are more strongly affected by seasonal changes than eukaryotic communities, reflecting their different adaptive strategies to environmental fluctuations (W. Huang et al., 2023). Additionally, it has been shown that eukaryotes positively influence prokaryotic diversity through multiple predation, enhancing complementarity and evenness among species (Saleem et al., 2012). Furthermore, various environmental conditions have been observed to influence the interaction strength between eukaryotic and prokaryotic organisms, leading to variations in microbial diversity, structure, and stability (Ohore et al., 2022). Therefore, understanding the interactions between eukaryotic and prokaryotic communities, as well as how these interactions vary under different environmental conditions, is essential for comprehensively understanding microbial community dynamics and their ecological implications.

These intricate interactions between eukaryotic and prokaryotic communities, along with the environmental pressures they face, are often reflected in the structure and dynamics of co-occurrence networks, which illuminate the mechanisms underlying the adaptability and resilience of microbial communities. These networks, which are constructed based on correlations between species abundances, provide insights into the structural properties of microbial communities, including their stability, robustness, and the potential for species to co-adapt or compete within their environments. Previous studies have indicated that higher biodiversity may influence the stability of co-occurrence networks, with weak interactions and co-exclusion contributing to the greater network of temporal stability and resilience, depending on the complexity of species interactions (Coyte et al., 2015). Modularity quantifies the extent to which taxa are grouped into interacting or coexisting units, known as modules. This topology of the network reflects various biological processes, such as shared ecological functions within modules, spatial segregation, or similar habitat preferences among taxa. Modularity can significantly impact community stability. For instance, high modularity stabilizes communities by containing the effects of a taxon’s loss within its module, preventing the disruption from spreading throughout the entire network. However, higher environmental pressures may lead to lower network modularity and an increase in positive associations, which could be key processes that disrupt community stability and ultimately destabilize microbial networks (Hernandez et al., 2021). This underscores the critical need for further research into the interplay between biodiversity, network structure, and environmental pressures to safeguard microbial community stability and ecosystem health

In this study, we focused on the short-term temporal dynamics of microbial communities, employing high-throughput amplicon sequencing of 16S rRNA and 18S rRNA genes to comprehensively analyze microbial diversity and community structure, allowing for continuous monitoring of both prokaryotic and eukaryotic communities. To explore the ecological relationships and network properties of these communities, we constructed co-occurrence networks based on Spearman correlations and evaluated their stability and robustness using simulations. Additionally, we employed partial least squares path modeling (PLS-PM) to assess the interactions between eukaryotic and prokaryotic communities and their impact on network complexity. This study provides a comprehensive understanding of how microbial communities in aquatic systems respond to short-term temporal variations, emphasizing the critical interplay between biodiversity, network structure, and ecosystem stability.

## 2 Material and Method

### 2.1 Sample collection and environmental attributes analysis

The Haihe River, a critical water resource in Tianjin, China, serves multiple functions including drainage, water storage, supply, shipping, tourism, and environmental protection. The study area is located at Lishunde Wharf (117.22° E, 39.12° N) on the Haihe River in Tianjin. This region is predominantly influenced by the East Asian monsoon and exhibits a temperate monsoon climate. Sampling was conducted over 88 days, from May 28, 2021, to August 23, 2021, with samples collected every three days, resulting in a total of 30 samples. In total, 10 L of in-situ surface water (about 0.5 cm depth) was collected using a plexiglass water collector. Surface water (500 mL) from each sampling site was immediately filtered using a 0.22 μm polycarbonate membrane (Millipore, USA). The samples were stored at low temperatures and transported to the laboratory as soon as possible for water quality analysis. The water quality parameters analyzed included total nitrogen (TN), total phosphorus (TP), ammonium nitrogen (NH_4_-N), nitrate nitrogen (NO_3_-N), total suspended solids (TSS), and pH.

TN was measured by oxidation with potassium persulfate, and absorbance was determined at 220 nm and 275 nm using a UV spectrophotometer. TP was digested with 5% potassium persulfate and measured using the molybdenum-antimony anti-spectrophotometric method at 700 nm. NH_4_-N was determined by the Nessler reagent method, with absorbance measured at 420 nm. NO_3_-N was measured using the zinc-cadmium reduction method, with absorbance determined at 543 nm. pH was measured by immersing the electrode of a pH meter in the water sample and recording the stabilized data.

### 2.2 Amplicon sequencing and processing

The microbial DNA was extracted using the Magen HiPure Soil DNA Kit (Magen, Guangzhou, China). The concentration of the extracted DNA was measured with a Qubit fluorometer (Thermo Scientific, USA), and its purity was assessed using a Nanodrop 2000 (Thermo Scientific, USA). The hypervariable region V3–V4 of the bacterial 16S rRNA gene was amplified using the primer pair 338F/806R, while the hypervariable region V4 of the eukaryotic 18S rRNA gene was amplified with the primer pair 528F/706R. The PCR products were first evaluated via 1% agarose electrophoresis and then purified using the QIAquick PCR Purification Kit (Qiagen, Germany). The purified PCR products were pooled in equal amounts and sequenced on the Illumina NovaSeq PE250 platform at Frasergen (Wuhan, China).

The raw FASTQ files were processed using QIIME2 v2023.2. Initially, the paired-end reads were merged and quality-filtered using VSEARCH with default parameters (Rognes et al., 2016). The denoising and generation of amplicon sequence variants (ASVs) were performed using the DADA2 pipeline (Callahan et al., 2016). Taxonomic classification of each ASV was carried out using the Silva database v138.1 (Quast et al., 2012). To ensure comparability across samples, microbial profiles were rarefied to the same sequencing depth using the “Rarefy” function in the “GuniFrac” package in R v4.2.1 (Chen et al., 2012). All subsequent analyses were conducted in R unless otherwise specified.

### 2.3 Statistical and Ecological Analysis

The α-diversity index (Shannon, Simpson, and Richness) for each sample and the β-diversity index (“Bray-Curtis” distance) for each pairwise sample both using the “vegan” package. The sum of the total phylogenetic branch length (phylogenetic distance, PD) was calculated using the “picante” package (Faith, 1992). To investigate the temporal dynamics of microbial diversity, we performed a quadratic polynomial regression analysis on the α-diversity and phylogenetic distance. Additionally, the time decay relationship (TDR) was analyzed using a quadratic polynomial regression to capture potential non-linear relationships. Microbial community stability was evaluated using the average variation degree (AVD) (Xun et al., 2021). The changes in species abundance over time were described using mean rank shifts (MRS), calculated with the “codyn” package (Hallett et al., 2016). To examine the relationships between physicochemical properties and the microbial community structure, Mantel tests were performed. The hierarchical partitioning (HP) was used to further analyze the significance of individual environmental factors and their combined effects by “rdacca.hp” package (Lai et al., 2022).

All co-occurrence network analysis was done using the ’meconetcomp’ package (Liu et al., 2023). Co-occurrence networks were constructed based on Spearman correlation coefficients. To ensure the robustness of the network, only species that appeared consistently for 12 consecutive days were included in the analysis. A correlation was considered robust when the absolute value of the correlation coefficient (|r|) was greater than 0.85 and the p-value was less than 0.01. The p-values were adjusted using the false discovery rate (FDR) method. To assess the stability and robustness of the networks under species extinction scenarios, simulations were conducted by randomly removing nodes in each network. The average degree and natural connectivity were then computed to evaluate network stability. We generated sub-networks for each time-series sample from meta-community networks by preserving ASVs presented in each time point sample. Network-level topological features provided in “subnet_property” function were calculated for each sub-network. The visualization was carried out using the Gephi v0.10.1 (Bastian et al., 2009)

We used partial least squares path modeling (PLS-PM) to evaluate the interactions between eukaryotic and prokaryotic communities and their impact on co-occurrence networks (Tenenhaus et al., 2005). The complexity of the network was characterized using latent variables constructed from Density, Average Degree, Average Path Length, and Complexity Heterogeneity. The goodness of fit (GoF) was used to assess the overall fit and robustness of the model.

## 3 Result

### 3.1 Dynamics of microbial community composition and diversity

We collected a series of time-series samples, where the V9 region of the eukaryotic small-subunit 18S rRNA gene and the V3-V4 region of the prokaryotic 16S rRNA gene were amplified and sequenced for comprehensive analysis of microbial community dynamics. Subsequently, the data was processed, which included quality filtered, denoised, merged, chimera removal, and rarefaction. The prokaryotes retained 34342 merged sequences per sample, which were clustered into 8439 ASVs, and the eukaryotes retained 47089 per sample, which were clustered into 5454 ASVs.

Subsequently, to estimate trends in community the α-diversity (Shannon, Simpson, and Richness) and phylogenetic distance over time, we fitted both the prokaryotic and eukaryotic data using a quadratic model that included time and its squared term to account for possible nonlinear temporal effects, the result in shown in Fig. 1A. For prokaryotes, the diversity metrics generally exhibit a nonlinear pattern, with initial increases followed by decreases over time. Specifically, the Shannon diversity shows an initial increase followed by a decrease (R² = 0.25, *p* _quadratic_ < 0.01, *p* _linear=_ 0.93). The Simpson diversity also demonstrates a nonlinear trend (R² = 0.32, *p* _linear_ = 0.85, *p* _quadratic_ < 0.01). Richness diversity similarly exhibits a nonlinear pattern (R² = 0.23, *p* _linear_ < 0.05, *p* _quadratic_ = 0.1). phylogenetic distance exhibits both linear and nonlinear changed (R² = 0.39, *p* _linear_ < 0.01, *p* _quadratic_ < 0.05). For eukaryotes, the diversity metrics show a consistent and significant increase over time. The Shannon diversity increases significantly (R² = 0.47, *p* _linear_ < 0.001, *p* _quadratic_ = 0.93). The Simpson diversity also increases significantly (R² = 0.41, *p* _linear_ < 0.001, *p* _quadratic_ = 0.99). Richness diversity exhibits a significant linear increase (R² = 0.45, *p* _linear_ < 0.001, *p* _quadratic_ = 0.91). Phylogenetic distance shows a moderate increase (R² = 0.17, *p* _linear_ < 0.05, *p* _quadratic_ = 0.43).

**Fig. 1.**
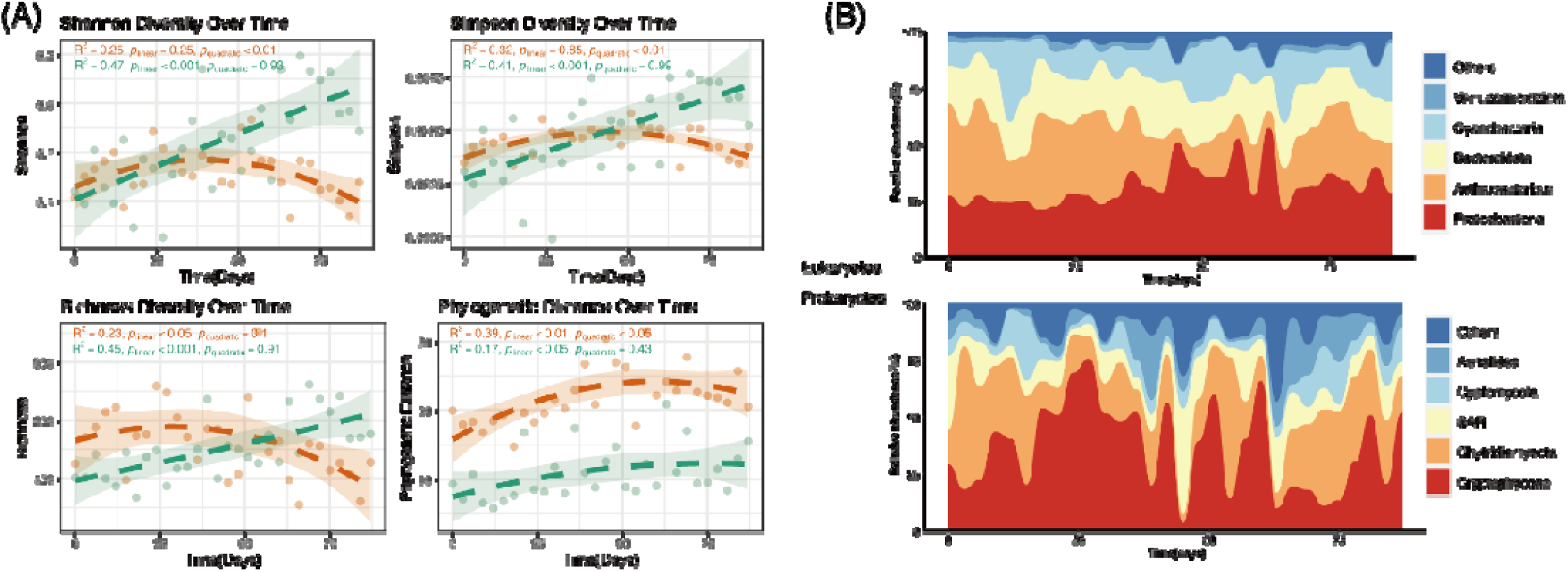
Temporal dynamics of microbial community diversity and composition. 834 **(A)** Temporal trends in microbial community diversity, including Shannon diversity, Simpson diversity, Richness, and Phylogenetic Distance for both eukaryotic and prokaryotic communities. **(B)** Temporal trends in the taxonomic composition of the microbial community for eukaryotes and prokaryotes.

The community composition of eukaryotes and prokaryotes was analyzed over time (Fig. 1B). The eukaryotic community was predominantly composed of Cryptophyceae (36.62%), Chytridiomycota (25.17%), and SAR (11.32%), while the prokaryotic community was primarily composed of Proteobacteria (31.39%), Actinobacteriota (25.83%), and Bacteroidota (20.33%). Variance analysis was used to evaluate fluctuations in the relative abundance of different phyla over time. Among prokaryotes, Actinobacteriota, Proteobacteria, and Cyanobacteria exhibited the highest variances. For eukaryotes, Cryptophyceae, Chytridiomycota, and SAR showed the greatest variances.

### 3.2 Dynamics of Microbial Community Structure and Environmental Influences

The time-decay relationships (TDRs) for microbial communities were modeled by fitting the Bray-Curtis dissimilarity of prokaryotic and eukaryotic communities using both time and its quadratic term (Fig. 2A). For prokaryotic community, the fitted quadratic model shows a significant linear increase in dissimilarity over time (*p* _linear_ < 0.001, *p* _quadratic_= 0.082), the distribution of Bray-Curtis dissimilarity show a higher density of values close to 1, indicating that prokaryotic community underwent a directional changed, more variation in community structure over time. In contrast, eukaryotic community dissimilarity shows a nonlinear trend with initial decreases followed by increases over time (*p* _linear_ = 0.47, *p* _quadratic_< 0.05), have a wider range of dissimilarity values, with a notable peak around 0.8, Eukaryotic communities have undergone a process of community homogenization followed by heterogenization, but in general, relatively minimal temporal changes in community structure. Meanwhile, we assessed the community stability of eukaryotic and prokaryotic communities across two phases using the average variation degree (AVD). The results indicated that during the heterogenization phase, community stability was significantly lower compared to the homogenization phase. Moreover, there was a significant difference (p < 0.01) in the stability between eukaryotic and prokaryotic communities (Fig. S1).

**Fig. 2.**
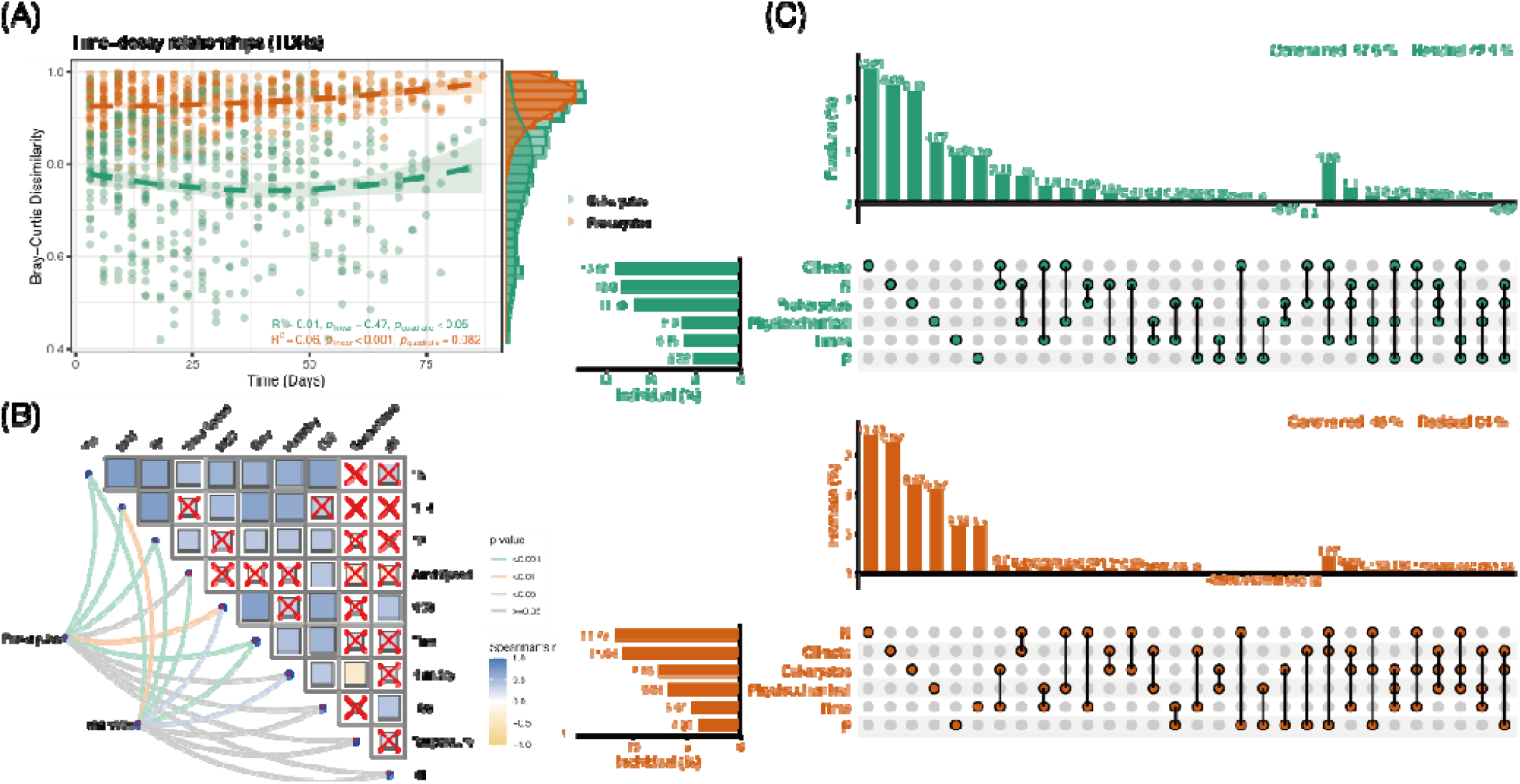
Temporal dynamics of microbial community structure and environmental influences. **(A)** Time-decay relationships (TDRs) for prokaryotic and eukaryotic communities, showing Bray-Curtis dissimilarity over time. **(B)** Mantel test results showing correlations between microbial community structure and environmental factors for both prokaryotes and eukaryotes. **(C)** Hierarchical partitioning analysis, showing the contribution of individual and combined environmental factors to the variation in microbial community structure for prokaryotic and eukaryotic communities.

Following, we conducted a Principal Coordinates Analysis (PCoA) based on the Bray-Crutis distance to further explore the temporal dynamics of microbial communities (Fig. S2). The analysis reveals a clear separation of samples along the primary coordinate axis, indicating distinct temporal changes in community structure. The ellipses represent the 95% confidence intervals for each time point group. The prokaryotic communities are distinctly separated over time, supporting the observed linear increase in Bray-Curtis dissimilarity, a clear separation is still evident along the primary coordinate axis. The eukaryotic communities show a more dispersed pattern, reflecting the nonlinear trend of initial decreases followed by increases in dissimilarity. Despite some overlap, the PERMANOVA results (F = 3.061, *p* = 0.001) confirm the statistical significance of these differences, the samples from different time points are distinguishable, indicating temporal shifts in community composition. In summary, the PCoA plots demonstrate that both prokaryotic and eukaryotic communities exhibit significant temporal changes in the middle of the time series. The prokaryotic communities show a more pronounced and consistent directional change, while the eukaryotic communities display a cyclical pattern of homogenization and heterogenization.

The unitary environmental drivers of the variation of microbial community were analyzed using Mantel test (Fig. 2B). The result revealed that the NH_4_^+^ had a significant impact on the prokaryotic community (*p* < 0.001), as well as TP and TN (*p* < 0.001). NO3 also significantly influenced the prokaryotic community structure (*p* < 0.01). For the eukaryotic community, TP and TN were significantly correlated with community structure (*p* < 0.001), along with Wind Speed (*p* < 0.001) and Time (*p* < 0.001). Among these environmental factors, TN exhibited the highest Mantel’s r value for the prokaryotic community, signifying it as the most influential factor. For the eukaryotic community, TP had the highest Mantel’s r value, indicating its strong association with community structure. Next, we employed hierarchical partitioning (HP) to address the contributions of multiple variables to community variation and their combined effects, effectively mitigating multicollinearity issues and identifying the most influential factors. For the eukaryotic community, factors collectively explained 57.5% of the variation, with climate factors having the strongest impact at 10.21%. The combined effects of all factors, including climate, prokaryotic community influences, and time, explained an additional 3.05% of the variation (Fig. 2C). For the prokaryotic community, the factors explained a total of 46% of the variation, with nitrogen (N) having the strongest impact at 10.46%. The combined effects of nitrogen, climate, and phosphorus (P) explained an additional 1.27% of the variation in the prokaryotic community structure (Fig. 2D).

### 3.3 Co-occurrence pattern of microbial community

To further elucidate the periods of community homogenization and heterogenization, we analyzed the microbial co-occurrence networks. The analysis was based on the Spearman correlation coefficients, which allowed us to generate comprehensive co-occurrence networks for both phases (Fig. 3A). The analysis revealed that both networks have a similar number of nodes, with the homogenization network comprising 722 nodes and the heterogenization network comprising 742 nodes. However, the proportion of prokaryotic nodes decreased significantly from 38.09% in the homogenization phase to 23.32% in the heterogenization phase. Notably, the heterogenization network has twice as many edges (6429) compared to the homogenization network (3209), resulting in an average degree of 17.33, almost double that of the homogenization network (8.89). The centralization metric, which measures the dominance of key nodes, is higher in the heterogenization network (0.098) than in the homogenization network (0.036), indicating a more prominent presence of key nodes or hubs in the heterogenization network. In contrast, the modularity is higher in the homogenization network (0.920) compared to the heterogenization network (0.746), suggesting a more distinct community structure in the homogenization network. Overall, from phase I to phase II, the heterogenization networks exhibit larger network diameters and increased complexity. These networks tend to become more centralized, with greater diversity in node connectivity, as indicated by high node heterogeneity. These findings highlight the transition from a less connected, modular structure in the homogenization phase to a more interconnected and centralized network in the heterogenization phase (Table S1).

**Fig. 3.**
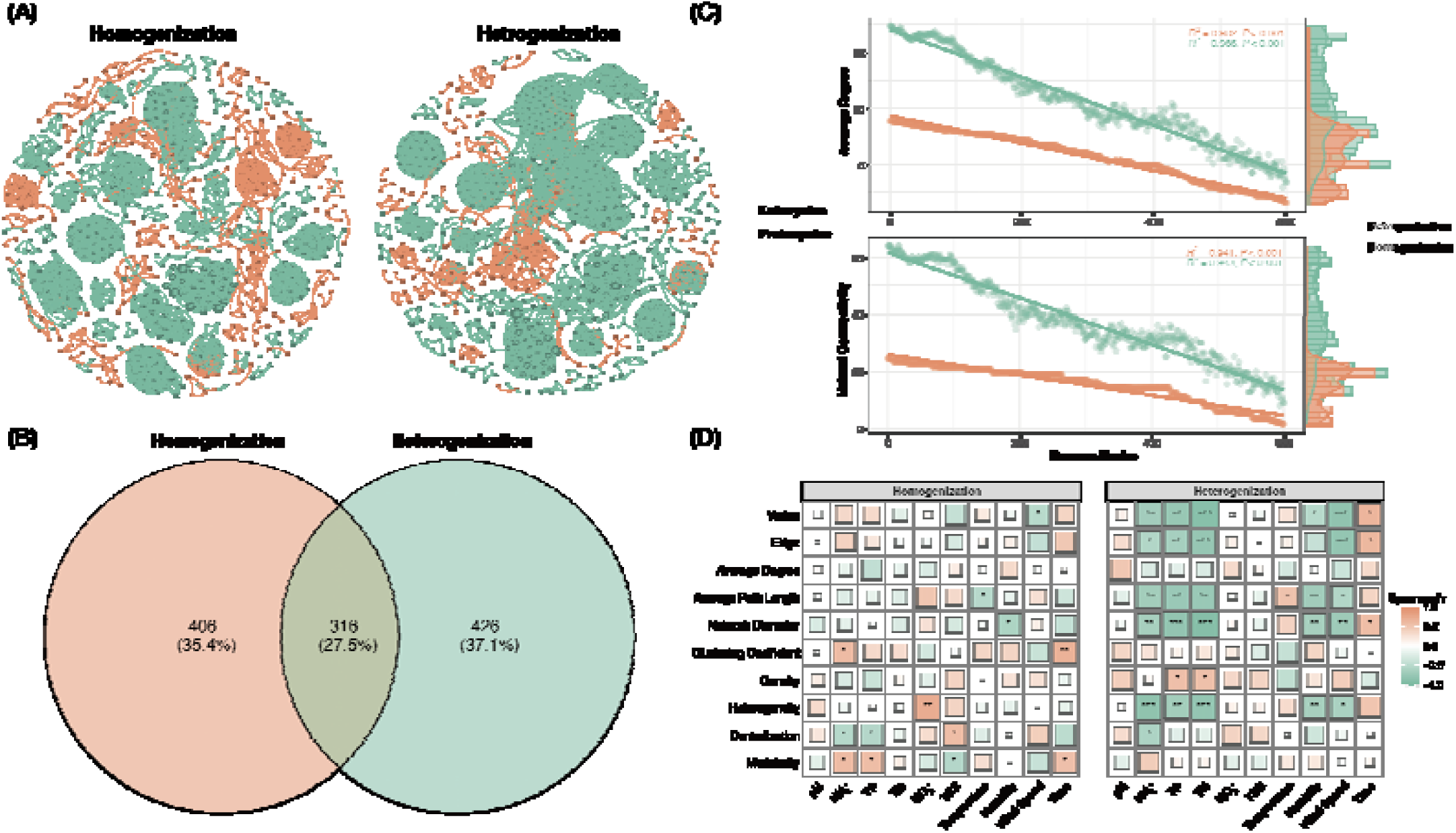
Co-occurrence network analysis of microbial communities during periods of homogenization and heterogenization. **(A)** The co-occurrence networks of microbial communities during the homogenization (left) and heterogenization (right) phases. Nodes represent different taxa, and edges represent significant correlations (Spearman correlation, |r| > 0.85, p < 0.01). The size of the nodes is proportional to their degree of connectivity. **(B)** The overlap of microbial taxa between the homogenization and heterogenization networks. **(C)** Network robustness analysis based on average degree and natural connectivity metrics, comparing the stability of homogenization and heterogenization networks under simulated node removal scenarios. **(D)** The relationships between environmental variables and network properties during the homogenization and heterogenization phases. Green indicates positive correlations, while red indicates negative correlations. The intensity of the color represents the strength of the correlation.

Subsequently, we analyzed the overlap of nodes between the two networks. The results showed that the two networks share 316 nodes, accounting for 27.5% of the total nodes in both networks. The homogenization network has 406 unique nodes, representing 35.4% of its total nodes, while the heterogenization network has 426 unique nodes, making up 37.1% of its total nodes (Fig. 3B). Among these unique nodes, the number of prokaryotic nodes decreased from 223 in the homogenization network to 121 in the heterogenization network. Conversely, the number of eukaryotic nodes increased from 183 in the homogenization network to 306 in the heterogenization network. To further investigate the diversity of node connectivity and the robustness of the two networks, we simulated species extinction by randomly removing nodes from the networks (Fig. 3C). The stability of the network was assessed using the average degree and the natural connectivity index. Although the average degree and natural connectivity of heterogeneous networks are higher than that of homogenization networks, the average degree and natural connectivity decrease faster in heterogenization networks than in homogenization networks. This suggests that the homogenization network is more stable to node loss, exhibiting stronger resistance and indicating higher network robustness. The scatter density function also shows that, in the initial stages of node removal, the average degree and natural connectivity in the homogenization network are more concentrated, demonstrating higher robustness. Conversely, the scatter density distribution in the heterogenization network is more dispersed, indicating that it is more prone to abrupt changes during the process of node removal. Removal of highly connected nodes (critical nodes) significantly affects the network connectivity. We further investigate the vulnerability of the nodes in both networks, as measured by each node’s network efficiency (Eff). The results show that the median node vulnerability in the homogenization network (0.009) was originally smaller than that in the heterogenization network (0.283), supporting the above results (Fig. S3).

We constructed subnetworks for each sample to examine the influence of environmental factors on network properties. Each subnetwork contains the nodes present in the respective sample, allowing us to extract and analyze the subnetworks (thereby calculating the attributes for each subnetwork). Spearman correlation was then used to assess the relationship between the subnetwork properties and environmental factors. The analysis revealed that environmental factors significantly influence network properties in both homogenization and heterogenization networks, with distinct patterns observed between the two phases. In the homogenization network, NH_4_^+^ showed a positive correlation with vertex count and edge number, while TP was correlated with network diameter and clustering coefficient. TN was associated with average path length and density. Temperature and wind speed also influenced various network properties. In the heterogenization network, NH_4_^+^ had strong correlations with vertex count, edge number, and average degree, while TP and TN were linked to multiple network attributes, highlighting their crucial role in shaping network structure. NO_3_^-^ was correlated with network diameter and average path length, indicating its role in network complexity. Time-affected clustering coefficient and centralization, reflecting temporal dynamics (Fig. 3D). Overall, heterogeneous networks have a stronger and broader correlation with environmental variables, which suggests that heterogeneous networks are more fragile and susceptible to environmental factors with higher complexity and dynamics than homogeneous networks.

### 3.4 Potential Drivers of the microbial community structure and network properties

We further explore the relationships between eukaryotic and prokaryotic communities, as well as the inter-domain co-occurrence network properties, using Partial Least Squares Path Modeling (PLS-PM). The PLS model’s goodness of fit (GoF) score of 0.60 indicates a good explanatory power, capturing significant patterns within the dataset (Fig. 4).

**Fig. 4.**
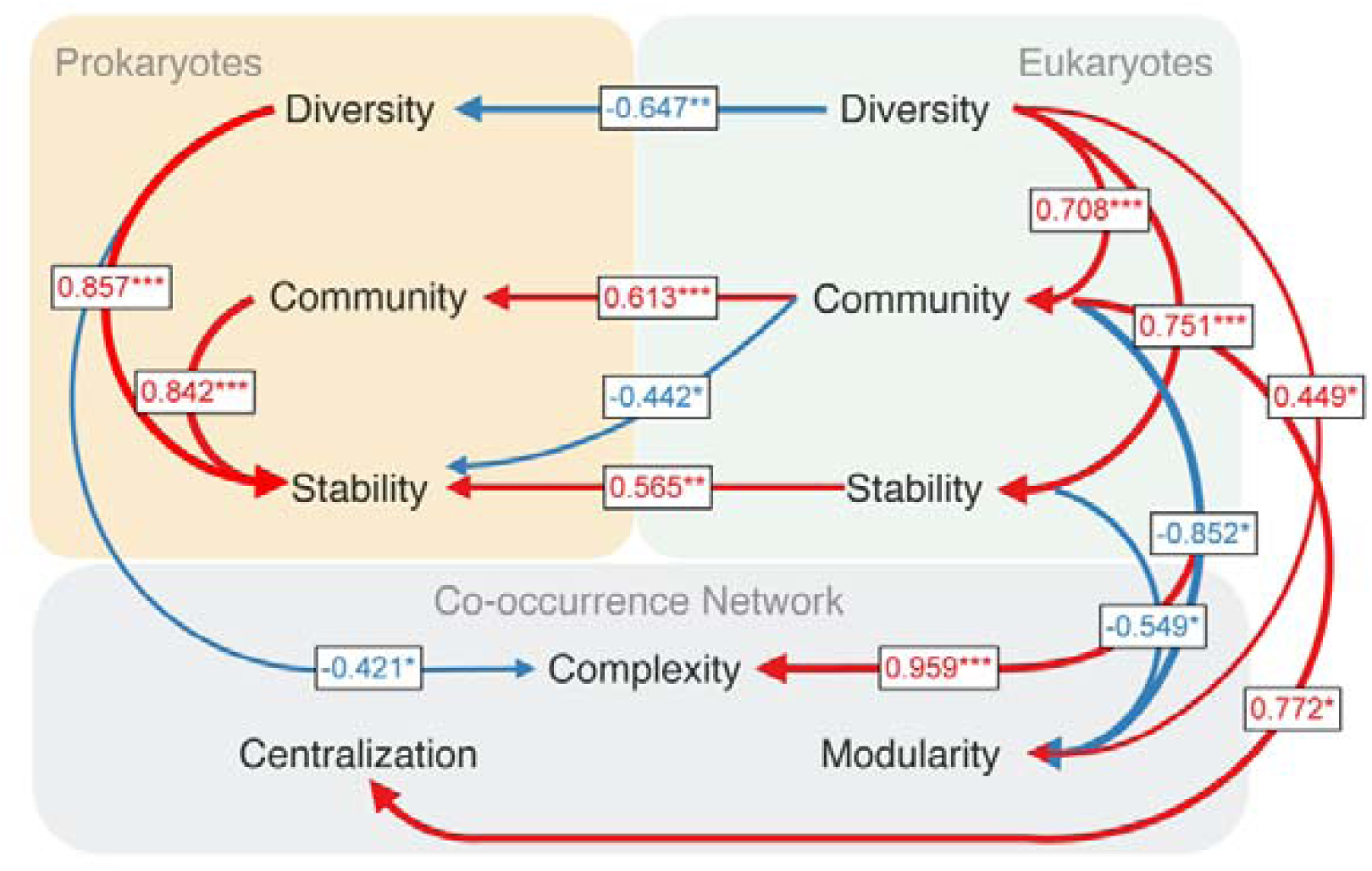
Structural equation modeling (SEM) illustrating the interactions between eukaryotic and prokaryotic communities and their influence on co-occurrence network properties. Arrows represent the direction of influence, with red indicating a positive effect and blue indicating a negative effect. Numbers on the arrows represent the standardized path coefficients, with asterisks indicating the level of statistical significance (*p < 0.05, **p < 0.01, ***p < 0.001).

The analysis reveals a strong positive relationship within the eukaryotic community. Eukaryotic community diversity strongly positively influences eukaryotic stability (coefficient = 0.751, *p* < 0.001) and eukaryotic community structure (coefficient = 0.708, *p* < 0.001). Similarly, prokaryotic community structure (coefficient = 0.842, *p* < 0.001) and diversity (coefficient = 0.857, p < 0.001) have a strongly positive effect on stability. Additionally, the eukaryotic community strong influence on the prokaryotic community, including a notable positive correlation of eukaryotic community structure on prokaryotic community structure (coefficient = 0.613, *p* < 0.001) and stability (coefficient = 0.565, *p* < 0.01). However, there are also significant negative relationships, such as between eukaryotic diversity and prokaryotic diversity (coefficient = -0.647, *p* < 0.01), and between eukaryotic community structure and prokaryotic stability (coefficient = -0.442, *p* < 0.05).

The model also sheds light on how characteristics of eukaryotic and prokaryotic communities influence network properties. The model also sheds light on how microbial community characteristics influence network properties. The eukaryotic community structure shows a strong positive association with network Complexity (coefficient = 0.959, *p* < 0.001) and Centralization (coefficient = 0.772, *p* < 0.05), indicating that more diverse and stable eukaryotic communities contribute to more complex and centralized network structures. Additionally, the eukaryotic community structure negatively impacts network Modularity (coefficient = -0.852, *p* < 0.05), suggesting that as the eukaryotic community becomes more interconnected, the network becomes less modular. On the prokaryotic side, diversity harms network Complexity (coefficient = -0.421, *p* < 0.05).

## 4 Discussion

### 4.1 Distinct Response Patterns in Eukaryotic and Prokaryotic Community Diversity

Our findings reveal distinct temporal response patterns of eukaryotic and prokaryotic microbial communities. Specifically, the alpha diversity of eukaryotic communities (including Shannon, Simpson, and richness diversity) and phylogenetic distance showed a significant linear increase throughout the study period. This consistent increase in diversity suggests that the eukaryotic communities experienced a continual enhancement in species diversity and evenness over time. The Bray-Curtis dissimilarity for eukaryotic communities displayed a nonlinear trend, with initial decreases followed by subsequent increases. The eukaryotic samples exhibited a more dispersed pattern in the PCoA, reflecting the initial homogenization and subsequent heterogenization trends observed in the Bray-Curtis dissimilarity analysis. Despite some overlap, the temporal changes in eukaryotic community composition were significant (PERMANOVA F = 3.061, p = 0.001). This pattern can be explained by the initial phase of community homogenization, likely driven by favorable and stable environmental conditions that provided high ecological redundancy, allowing the establishment and growth of a wide range of species (Feng et al., 2018). As time progressed and environmental conditions began to fluctuate or resources became more limited, the ecological redundancy decreased. This resulted in increased competition and differentiation among species, leading to a reorganization of the community structure and a subsequent increase in dissimilarity (Bloxham et al., 2024). This non-linear dynamic highlights the adaptive and resilient nature of eukaryotic microbial communities in response to environmental fluctuations (C. Guo et al., 2022). During the study period, the favorable environmental conditions may have facilitated the introduction and establishment of a broader range of eukaryotic species. The availability and diversity of resources could have provided niches for different species, promoting higher species richness and evenness initially. As environmental conditions changed, possibly due to seasonal variations, the eukaryotic communities underwent structural reorganization. This led to differentiation among the microbial populations, resulting in an increase in dissimilarity. Mantel test and hierarchical partitioning analyses indicated that climate factors, such as wind speed, significantly influenced eukaryotic community structure (p < 0.001). These factors likely played a pivotal role in driving the observed increase in diversity by affecting resource distribution and creating microhabitats that favored different species over time. Overall, these phenomena suggest that increased community diversity fosters a non-directional adaptive capacity and resilience to environmental conditions, underscoring the critical role of diversity in enhancing functional redundancy and ecological stability (Allison and Martiny, 2008).

In contrast, the prokaryotic microbial communities displayed greater fluctuations and sensitivity to environmental changes, presenting a markedly different and more passive pattern of quickly adaptive response. The Bray-Curtis dissimilarity for prokaryotic communities significantly increased (p < 0.001), indicating a directional change in community structure. The PCoA results showed significant separation of samples over time, corroborating the linear increase in Bray-Curtis dissimilarity. At the same time, the α-diversity of prokaryotic microbial communities exhibited a nonlinear pattern, initially increasing and then decreasing. This pattern indicates that during the early phase of the study, the introduction and establishment of new species increased diversity. However, as time progressed, competitive or environmental pressures likely led to the exclusion of some species, resulting in a decrease in diversity. The directional change of community suggests that the communities had high variability, driven by selective pressures such as environmental factors or interactions with eukaryotic communities, and exhibited strong responsiveness to these pressures (Nguyen et al., 2021). Under pressure, the species composition of prokaryotic communities rapidly changed, with species better suited to the environment achieving dominance. This resulted in significant shifts in community abundance and extensive reorganization of community structure, with less adapted species being excluded, leading to a decrease in diversity. This adaptive and resilient response pattern, driven by selective pressures, showcases the dramatic changes in community diversity and abundance. The high variability and responsiveness of prokaryotic communities to environmental changes suggest their ability to swiftly adapt to new conditions. Prokaryotes, with their shorter generation times and ability to engage in horizontal gene transfer, can quickly adjust their metabolic pathways to exploit changing resources and environmental niches (Philippot et al., 2021). Environmental factors, such as nutrient levels, temperature variations, and other physicochemical parameters, significantly influenced the prokaryotic community structure. Mantel test and hierarchical partitioning analyses indicated that these factors played a pivotal role in driving the observed increase in diversity by affecting resource distribution and creating microhabitats.

### 4.2 Network Centralization from Homogenization to Heterogenization

We further explored the differences in microbial co-occurrence networks during the periods of community homogenization and heterogenization. Using Spearman correlation coefficients, we generated comprehensive co-occurrence networks for both phases, providing deeper insights into the structural dynamics of prokaryotic-eukaryotic communities. The analysis revealed several key differences between the two networks. Both the homogenization and heterogenization networks have a similar number of nodes, with 722 and 742 nodes, respectively. However, the proportion of prokaryotic nodes decreased significantly from 38.09% in the homogenization phase to 23.32% in the heterogenization phase. This shift indicates that prokaryotic communities were more affected by the environmental changes, likely due to more diverse selection pressures during the heterogenization stage, which made certain prokaryotes unable to adapt and led to their exclusion. In contrast, the proportion of eukaryotic nodes increased significantly, suggesting that eukaryotic organisms could exploit diverse resources and ecological niches, showing greater adaptability and competitiveness during this stage (D. Huang et al., 2023; Prodinger et al., 2021). Notably, the heterogenization network had twice as many edges (6429) compared to the homogenization network (3209), resulting in a higher average degree of 17.33 versus 8.89. This increase in connectivity suggests a more complex and integrated network structure during the heterogenization phase. The increased environmental heterogeneity during this phase may have facilitated more complex species interactions and resource utilization, leading to tightly connected networks. Additionally, the centralization metric, which measures the dominance of key nodes, was higher in the heterogenization network (0.098) than in the homogenization network (0.036), indicating that certain keystone or “pivotal” species became more dominant in the network structure during the heterogenization phase. In contrast, the modularity was higher in the homogenization network (0.920) compared to the heterogenization network (0.746), suggesting a more distinct and compartmentalized community structure in the homogenization phase, with clearer boundaries between different microbial subgroups. The stable and uniform environmental conditions during the homogenization phase likely contributed to this more segmented and independent community structure.

Ecological interaction networks exhibit a coherent structure characterized by well-defined patterns that contribute to network stability. Network stability is influenced by species connectivity, the presence of negative interactions, and a balance of a few strong and many weak interactions (Coyte et al., 2015). Additionally, the clustering of species into subgroups, known as modularity, involves numerous connections within species subgroups and fewer connections between them, effectively decoupling the subgroups within the community. This modularity minimizes the risk of changes during perturbations by creating dependencies between species, promoting species asymmetry, and buffering against the propagation of perturbation effects across different parts of the network (Garay□Narváez et al., 2014). On the other hand, centralization, which measures the dominance of key nodes, can lead to a network where certain keystone or “pivotal” species become more influential. While centralization can enhance network efficiency and robustness by facilitating quick responses to changes, it also makes the network more vulnerable to the loss of these key species, potentially leading to significant disruptions in network stability (Wu et al., 2019). We observed similar findings in our study. To investigate the diversity of nodes and network robustness, we simulated species extinction by randomly removing nodes. The results showed that although heterogeneous networks have higher connectivity, these metrics decrease faster compared to homogenization networks. This suggests that homogenization networks are more stable and resistant to node loss. The scatter density function further supports this, showing higher robustness in the homogenization network, whereas the heterogenization network is more prone to abrupt changes. We found similar observations by constructing subnetworks for each sample and using Spearman correlation analysis to examine the influence of environmental factors on network properties in both homogenization and heterogenization networks. The heterogeneous networks showed stronger and broader correlations with environmental variables, suggesting that they are more fragile and susceptible to environmental factors with higher complexity and dynamics compared to homogeneous networks.

### 4.3 Eukaryotic Fluctuations Drive Integrate Network Centralization

The relationships between eukaryotic and prokaryotic communities, as well as their inter-domain co-occurrence network properties, were explored using Partial Least Squares Path Modeling (PLS-PM), with the model achieving a GoF score of 0.60, indicating strong explanatory power. Previous research has shown that significant interactions exist between eukaryotic and prokaryotic communities, including important interactions such as predation and competition (He et al., 2023; Massana and Logares, 2013). Recent studies have also hinted that dissolved organic matter (DOM) may play a role in mediating the interactions between eukaryotic and prokaryotic communities, potentially influencing their ecological dynamics (Guan et al., 2024a, 2024b). Our results indicate that eukaryotic communities play a dominant role in shaping the interactions between eukaryotic and prokaryotic communities. This influence is multifaceted, affecting prokaryotic diversity, community structure, and stability. One of the significant findings is the negative relationship between eukaryotic diversity and prokaryotic diversity (coefficient = -0.647, p < 0.01). This suggests that higher eukaryotic diversity may lead to increased competition for resources, thereby reducing the diversity of prokaryotic communities. Eukaryotes, with their complex ecological roles and higher trophic status, can outcompete prokaryotes for limited nutrients and space, leading to a decline in prokaryotic diversity (Seymour et al., 2017). Eukaryotic community structure has a notable positive correlation with prokaryotic community structure (coefficient = 0.613, p < 0.001) and stability (coefficient = 0.565, p < 0.01). This indicates that changes in the eukaryotic community structure can lead to corresponding changes in the prokaryotic community structure. When eukaryotic communities are well-structured, they can create stable environments that promote the coexistence of various prokaryotic species. Conversely, significant structural changes in eukaryotic communities can disrupt prokaryotic communities, either through direct interactions or by altering the habitat (Huo et al., 2021; In ‘T Zandt et al., 2023).

Meanwhile, our model sheds light on how the characteristics of eukaryotic and prokaryotic communities influence co-occurrence network properties. The eukaryotic community structure shows a strong positive association with network complexity (coefficient = 0.959, p < 0.001) and centralization (coefficient = 0.772, p < 0.05). Additionally, the eukaryotic community structure negatively impacts network Modularity (coefficient = -0.852, *p* < 0.05). This phenomenon can be caused by several reasons. First, environmental stressors, such as pollution, nutrient limitation, or climate change, can induce niche compression, reducing the available resources and habitats within a community. Consequently, only species that are best adapted to the novel environmental conditions can survive and reproduce. These species often emerge as keystone or hub species, assuming more central and influential roles within the network, leading to network centralization (Banerjee et al., 2018). Additionally, as environmental pressures intensify, so too does intra-community competition. To maximize resource utilization and survival prospects, certain species gain a competitive advantage, dominating others through predation, competition, and symbiosis, further solidifying their central position in the community. As these keystone species become more central, the network’s centralized nature becomes more pronounced (Zeng et al., 2021). Furthermore, in response to environmental pressures, eukaryotic communities may reorganize their ecological networks to optimize function. For instance, by reducing redundant interactions or functions and strengthening key interactions, communities can improve resource use efficiency and overall resilience. This reorganization process can result in certain nodes (i.e., species) becoming more central and critical, thereby contributing to network centralization (Wang et al., 2021). Different species employ varying strategies to cope with environmental stressors. Some may enhance mutualistic and symbiotic relationships to improve survival chances, while others may adopt competitive and exclusionary tactics to maintain their niches (Sun et al., 2024). Overall, the dynamics of eukaryotic communities reflect this diversity of coping strategies, which can influence network structure and drive network centralization. While centralized networks may enhance functional stability in the short term due to the stabilizing effects of keystone species, excessive centralization can increase vulnerability to the loss of these critical species. Therefore, maintaining moderate diversity and network modularity is crucial for long-term ecological stability. In summary, eukaryotic communities often respond to environmental stressors by centralizing their networks to optimize survival and resource utilization. This process involves niche compression, competitive pressures, ecological network reorganization, diversity of coping strategies, and the trade-offs between biodiversity and stability. These factors collectively contribute to the centralization phenomenon observed in co-occurrence networks.

On the contrary, eukaryotic diversity shows a positive association with modularity (coefficient =0.449, p < 0.05), while the negative impact of prokaryotic diversity on complexity may also indirectly lead to an increase in network modularity (coefficient = -0.421, p < 0.05). This phenomenon can be caused by several reasons. Under favorable environmental conditions, increased diversity of eukaryotic and prokaryotic organisms promotes network modularity. This phenomenon can be attributed to the optimization of resource utilization and ecological function through modularity in biological communities. When resources are abundant and evenly distributed, more species can occupy distinct ecological niches, reducing direct competition. This niche differentiation fosters complementarity among species, enabling them to specialize in specific sub-environments and form modular structures (Li et al., 2022). Moreover, higher diversity is often associated with community stability and resilience. The complex interactions among species in diverse communities lead to the formation of multiple functional sub-groups or modules within the network. These modules can respond to environmental changes relatively independently, enhancing the overall stability and resilience of the ecosystem. Modular networks exhibit efficient local interactions and less efficient global interactions, improving overall system function. Under favorable conditions, species within communities can form tightly connected sub-groups, optimizing local resource utilization and energy flow. This optimization process contributes to the formation of modular structures, making the network more robust (Fang et al., 2023). Furthermore, favorable conditions promote the development of mutualistic relationships. These cooperative interactions can be optimized through the formation of modules, such as through nutrient complementarity, spatial utilization, and negative control. The strengthening of mutualistic relationships further promotes network modularity, as modules can effectively self-maintain and regulate. In favorable environments, biological communities enhance their adaptability and ecological redundancy through modular structures. Modular networks can accommodate more species and functions, and when one module is disrupted, other modules can continue to maintain ecosystem functions, thereby increasing the overall adaptability and resilience of the system (Y. Guo et al., 2022). Modular network structures help to limit the spread of disturbances, preventing local perturbations from rapidly cascading through the entire network. This characteristic is especially important in favorable environments as it allows ecosystems to remain stable despite local changes. By enhancing local interactions and partitioning, modular structures further improve ecosystem resilience and stability (Tian et al., 2017). In conclusion, under favorable conditions, the increased diversity of eukaryotic and prokaryotic organisms promotes network modularity. This process involves resource abundance, niche differentiation, community stability, increased diversity, mutualistic relationships, adaptability, and ecological redundancy. Modular structures optimize resource utilization, enhance functional efficiency, and improve ecosystem resilience, enabling biological communities to achieve optimal ecological function and stability in favorable environments.

## 5 Conclusions

In this study, we comprehensively analyzed the short-term temporal dynamics of microbial communities, revealing distinct response patterns between eukaryotic and prokaryotic communities. Our findings underscore the significant impact of environmental factors on microbial diversity and community structure, with eukaryotic communities showing gradual increases in diversity and stability, while prokaryotic communities exhibited more rapid and directional changes. The co-occurrence network analysis further demonstrated the critical role of network structure in mediating community responses to environmental fluctuations, highlighting the importance of modularity and centralization in maintaining ecosystem stability. These results contribute to our understanding of the adaptive mechanisms underlying microbial community dynamics, providing valuable insights into how biodiversity and network properties interact to sustain ecosystem functions.

## Supporting information

Graphical abstract

## Acknowledgement

This research was financially supported by the National Natural Science Foundation of China (NO. 32361133561) and the China Postdoctoral Science Foundation (2021M703430). Work of D.V.T. was supported by the Russian Science Foundation (grant no. 24-44-00093) https://rscf.ru/project/24-44-00093/

## Data availability statement

All the raw sequencing data were deposited into the NCBI Sequence Read Archive (SRA) database under accession number:

