## Supplementary figures and images for "Temporal Dynamics and Adaptive Mechanisms of Microbial Communities: Divergent Responses and Network Interactions"

### Graphical abstract

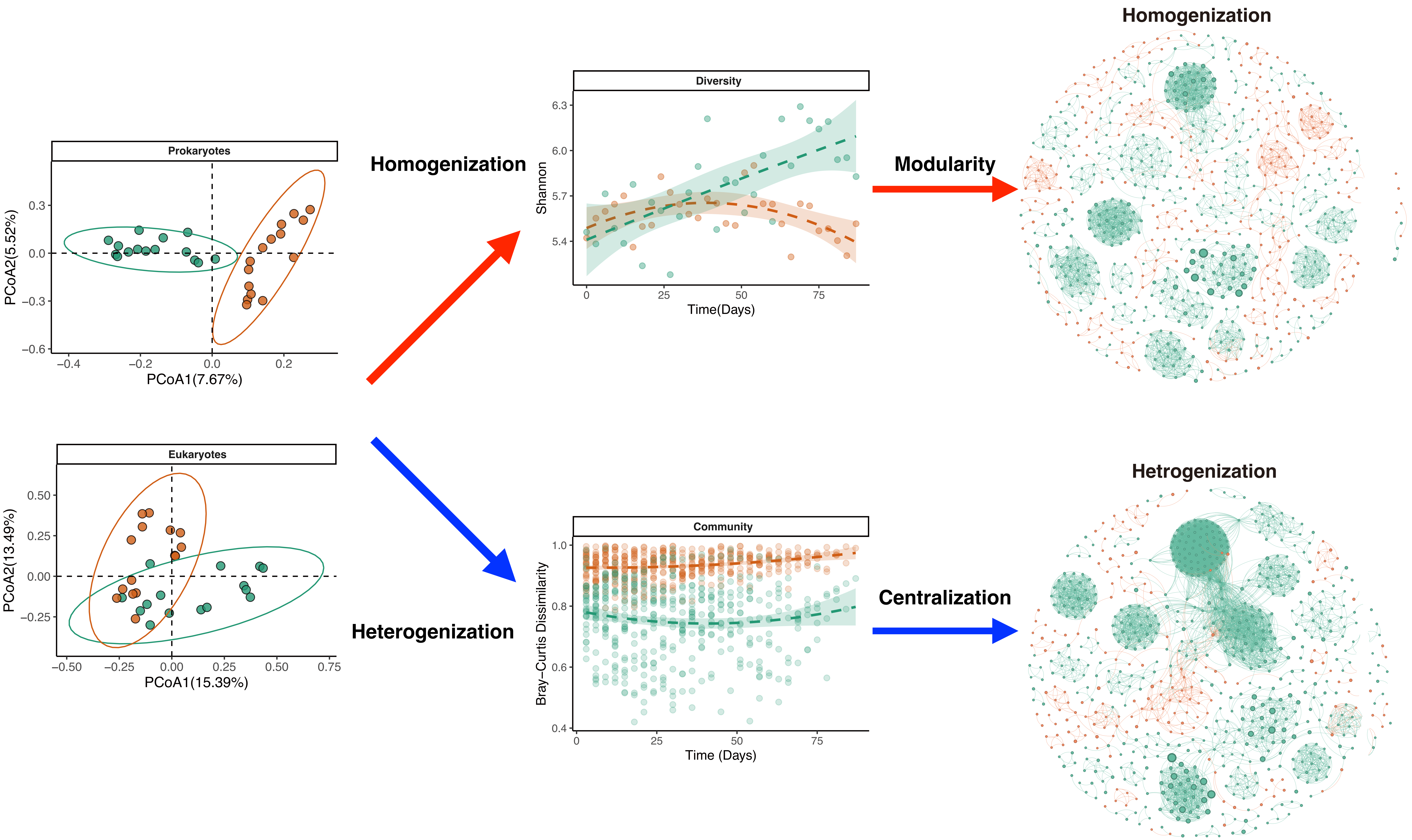

### Supplemental Figure 1

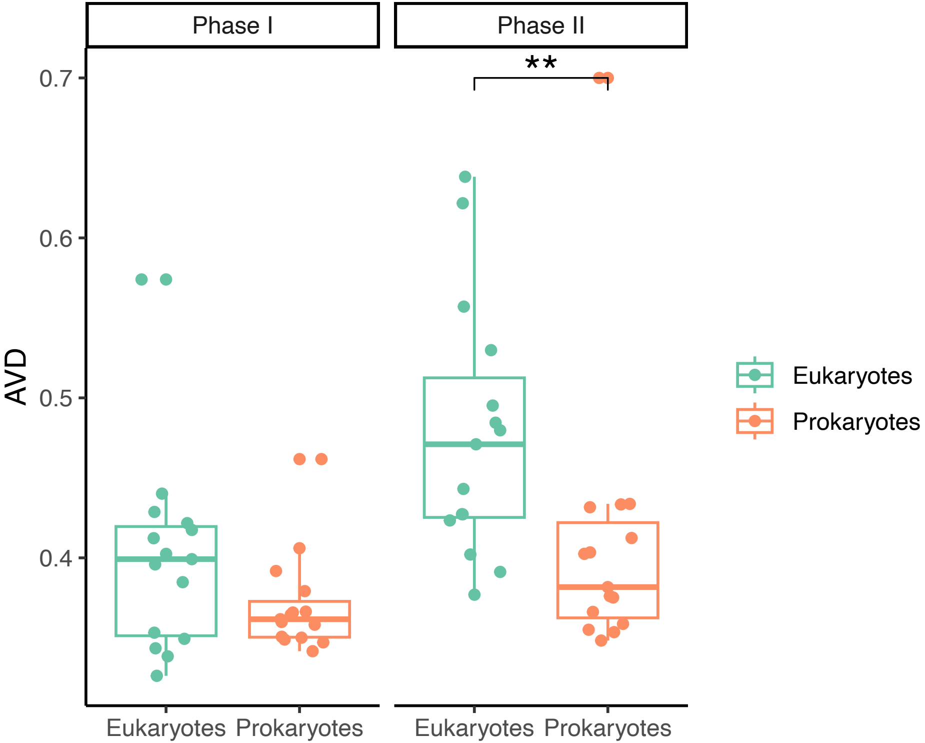

### Supplemental Figure 2

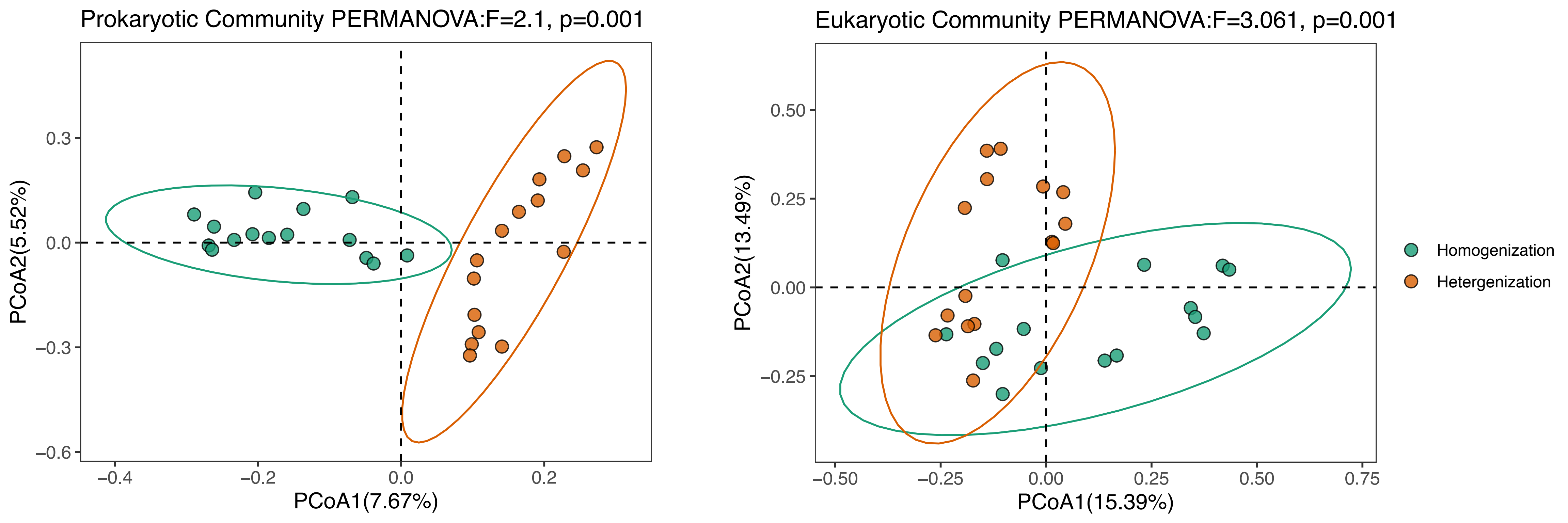

### Supplemental Figure 3

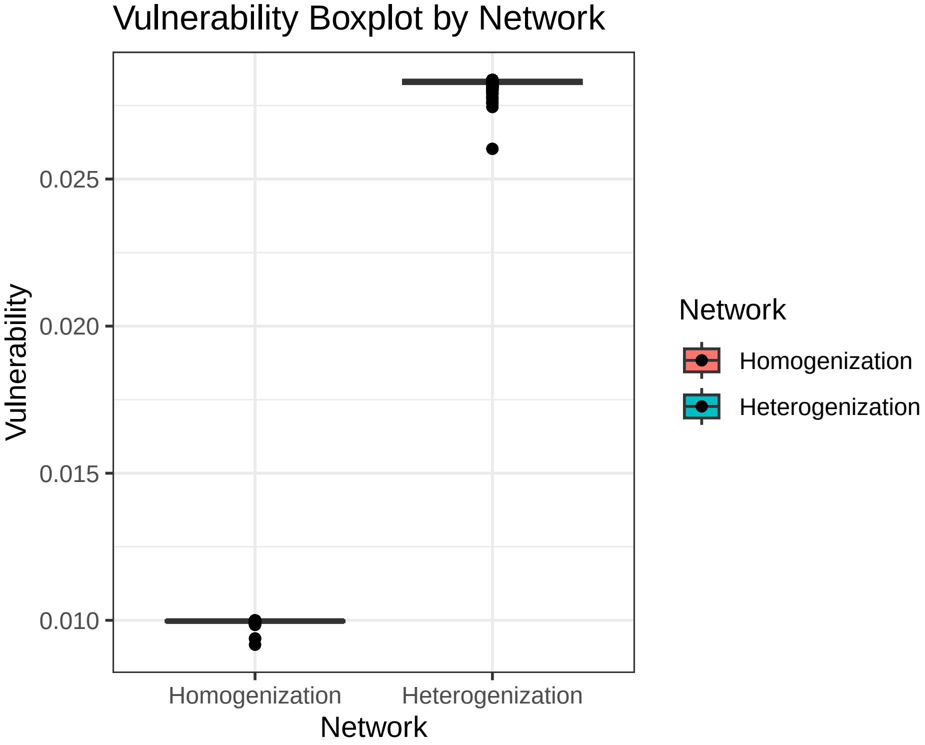
